# IPLS-LDA: An Improved Partial Least Square Discriminant Analysis for Heterogeneous Transcriptomics and Metabolomics Data Analysis

**DOI:** 10.1101/2022.11.02.514959

**Authors:** Snigdha Sarkar, Md. Shahjaman, Sukanta Das

## Abstract

Supervised machine learning (SML) is an approach that learns from training data with known category membership to predict the unlabeled test data. There are many SML approaches in the literature and most of them use a linear score to learn its classifier. However, these approaches fail to elucidate biodiversity from heterogeneous biomedical data. Therefore, their prediction accuracies become low. Partial Least Square Linear Discriminant Analysis (PLS-LDA) is widely used in gene expression (GE) and metabolomics datasets for predicting unlabelled test data. Nevertheless, it also does not consider the non-linearity and heterogeneity pattern of the datasets. Hence, in this study, an improved PLS-LDA (IPLS-LDA) was developed by capturing the heterogeneity of datasets through an unsupervised hierarchical clustering approach. In our approach a non-linear score was calculated by combining all the linear scores obtained from the clustering method. The performance of IPLS-LDA was investigated in a comparison with six frequently used SML methods (SVM, LDA, KNN, Naïve Bayes, RF, PLS-LDA) using one simulation data, one colon cancer gene expression data (GED) and one lung cancer metabolomics datasets. The resultant IPLS-LDA predictor achieved accuracy 0.841 using 10-fold cross validation in colon cancer data and accuracy 0.727 from two independent metabolomics data analysis. In both the cases IPLS-LDA outperformed other SML predictors. The proposed algorithm has been implemented in an R package, Uplsda was given in the https://github.com/snotjanu/UplsLda.

## 1. Introduction

Current world faced a great problem for unsorted, unformed and first shifting nature of large data. Most of the common large datasets in biological fields are OMICS. The gene expression datasets (GEDs) and metabolomics datasets are both the high-dimensional with fast progressive in nature that declares the facts from two separate biological criteria in the field of transcriptome and metabolome. A single gene expression data includes thousands of features that consists of small amount of samples with some heterogeneous feature that might be interrelated [1]. Therefore, the analysis of these complicated datasets becomes challenging in current situation [2,3]. On the other hand, there are two types of statistical machine learning: (a) supervised and (b) unsupervised learning methods which play important role in OMICS datasets. Classification is a supervised machine learning (SML) approach that accurately assigns unlabeled test data into specific categories. (e.g. normal or cancer) based on labeled training data [4].

Numerous SML techniques have been developed for mining GED and metabolomics data [5–11]. The earliest and widely used method is Fisher’s linear discriminant analysis (LDA) [12]. The co-linearity and curse of dimensionality is key drawbacks of LDA. K-Nearest Neighbor (KNN) [13] works by searching most nearest neighbor using the distance metric. In the Bayesian platform Naïve Bayes Classifier [14] is very popular classifier. Support vector machine (SVM) [15] has been extensively employed for analyzing different types of biomedical data. Random forests (RF) is an ensemble SML approach, it works by constructing a multitude of decision trees at training phase [16]. Partial least square (PLS)-regression has overcome the curse of dimensionality and multicollinearity problems of the OMICS data by transforming the input variable into the latent variable. The most popular variant of PLS-regression is Partial least squares discriminant Analysis (PLS-LDA) [17–20]. PLS-LDA encompassed with PLS-regression and LDA.

However, most of the classification techniques use linear scores for prediction of samples because of its simplicity and understandability. On the other hand, unsupervised classification techniques such as clustering have been successfully employed in unlabeled GED to elucidate biologically meaningful gene group on the basis of samples. They also play an important role for the discovery of new subtypes of diseases on the basis of genes [21]. The other applications of clustering in GED can be found in [22–28]. There are also many studies that combined both supervised and unsupervised classification methods. Shi et al. (2011) [29] used a low density separation (LDS) method for cancer classification. To discover the subtype specific patterns of the gene Wang et al. (2005) [30] used clustering methods. Then they calculated a score for each subtype. However, the methods that combined supervised and unsupervised classification as discussed earlier use linear scores and these methods are not convenient for real life applications because heterogeneity of the gene expression was not directly reflected in their prediction form. Hence, in this study, an improved PLS-LDA (IPLS-LDA) was developed by capturing the heterogeneity of datasets through an unsupervised hierarchical clustering approach. In our approach a non-linear score was calculated by combining all the linear scores obtained from the clustering method. Herein, we demonstrate the performance of the proposed classifier in a comparison of six popular classifiers such as SVM, LDA, KNN, naive Bayes classifier (NBC), random forest (RF) and PLS-LDA using one simulation data, one colon cancer gene expression data (GED) and one lung cancer metabolomics datasets. Results obtained from both types of data analysis confirm that the performance of the proposed method improve than the other popular classifiers.

In the next section, we described formation of the proposed IPLS-LDA classifier. In Section 3, we explore the real and simulation data analysis results of the proposed method in a comparison of some popular state-of-art methods. Finally, the paper will end in a conclusion.

## 2. Materials and Methods

### 2.1. Improved Partial least squares linear discriminant Analysis (PLS-LDA)

PLS-DA (Partial least square discriminant analysis) is the derivation of PLS-R (Partial least square regression) where the predictor vector Y contains discrete values. Let, *X* = (*x_ijk_*) be the ***n*** × ***p*** dimensional matrix where *x_ijk_* represents the expression of i^th^ samples and j^th^ gene with cluster *k* (*i= 1,2*, … … ….., *n*; *j=1,23*, … ….., *p_k_*; 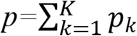). The equation of multiple linear regression model (MLR) can be expressed as:

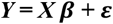

Where ***β*** is the regression coefficient of ***p*** × ***1*** matrix and the error vector ***ε*** is with ***n*** × ***1***. So ***Y*** is the ***n*** × ***1*** predictor variable vector. In this following approach, the least square solution is given by

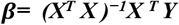

One of the major problems arises here when the number of gene is larger than the number of samples then ***X^T^X*** matrix becomes singular. Another complexity is that since gene expression datasets are the high dimensional and big datasets so many genes (variables) may be interrelated with each other therefore multicollinearity problem also arises here. Both PLS-DA and PLS-R solve this problem by decomposing the data matrix *X* into *m* (< *p*) orthogonal scores *U* (*n ×m*) with loadings matrix *Λ* (*p ×m*), and the predictor vector *Y*into *m* orthogonal scores *U* (*n × m*) and loadings matrix *λ* (*1 × m*). Let. *R* and *S* be the *n × p* and *n × 1* error matrices related with the data matrix *X* and predictor vector *Y* respectively. There are mainly two basic equations in the PLS-DA model

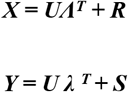

Now, consider a weight matrix *W* with *p ×m (orthogonal scores*), the scores matrix can be expressed

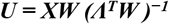

as and substituting it with the PLS-DA model,

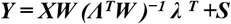

where 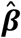, the regression coefficient vector is written as

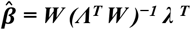

So the unknown sample value of Y can be predicted by,

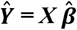

On simplification,

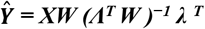

Linear scores obtained from PLS-LDA then combined by Kolmogorov-Nagumo average [1,31]

The PLS-DA algorithm estimates the matrices ***W, U, Λ***, and ***λ*** through the following steps:

#### Algorithm

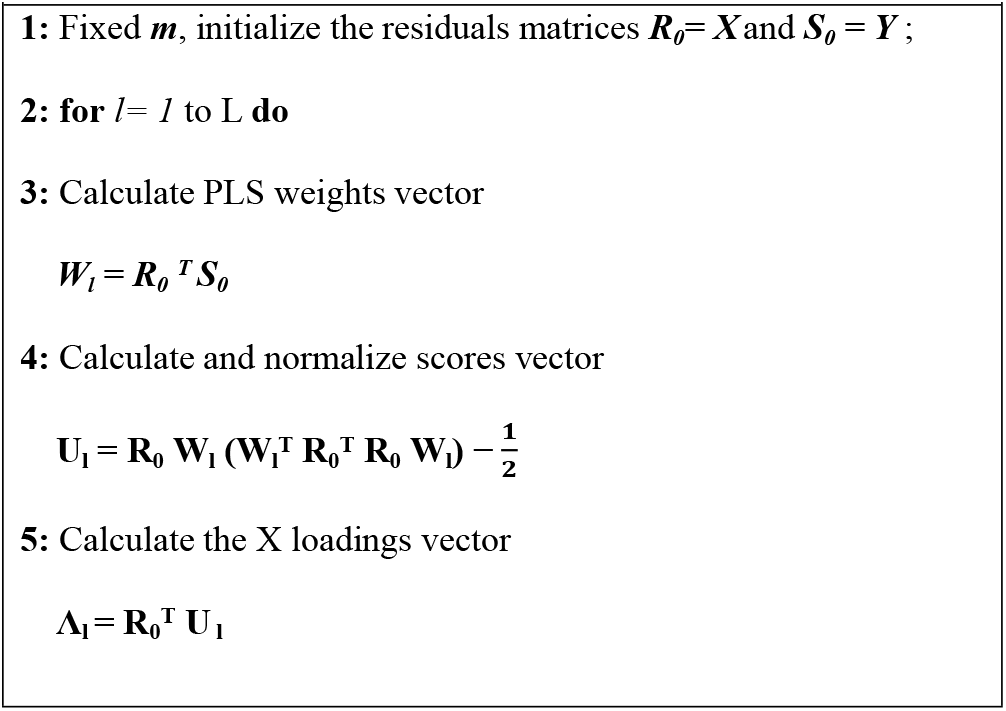
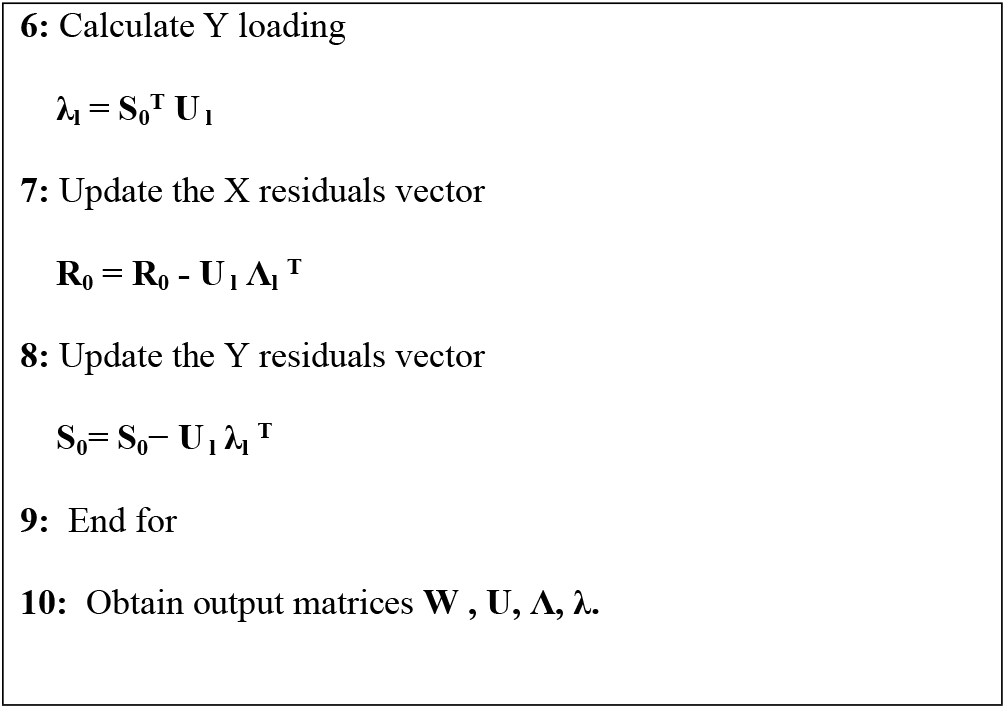

### 2.2. Performance Measures

To investigate the performance of different SML algorithms for binary sample classification with two groups such as normal and cancer, receiving operating characteristics (ROC) curve and area under the ROC curve (AUC) was used. We consider the following measures of performance:

False positive rate (FPR) = FP / (FP + TN),
Accuracy = (TP + TN) / (TP + FP + TN + FN),
Sensitivity = TP / (TP+ FN),
Specificity= TN / (TN +FN),
Positive predicted value (PPV)= TP / (TP+ FP),
Negative predicted value (NPV) = TN / (TN+ FN)
And Detection Rate = TP / (TP + FP + TN + FN)

Where, TP, TN, FP, FN, PPV, NPV denote the number of true positives, number of true negatives, the number of false-positive, number of false negatives, positive predicted values, negative predicted values respectively. A classifier is considered as best performer if it produces greater values of accuracy, sensitivity, specificity and lower values of FPR and NPV.

## 3. Results and Discussion

In this section, the performance of the proposed algorithm (**IPLS-LDA**) has been investigated in a comparison of six popular extensively used state-of-the-art methods (SVM, KNN, LDA, Naïve Bayes, Random Forest (RF) and PLS-LDA). Each of the algorithms was employed in a simulation dataset, a real colon cancer gene expression dataset and a metabolic dataset to check their performance. The performance measures of each method were calculated and recorded by the ROC package using R software. The other R packages were used in this study for the implementation of six competing classifiers are: plsgenomics, MASS, caret, e1071, random Forest, class and knn. All the R packages are available in the comprehensive R archive network (cran) or Bio-conductor.

### 3.1. Simulation data

The simulation data usually performed to mimic the characteristics of real gene expression data, which is shown in **Table 1** with two groups (*k*=2). Since real GED consists with gene corresponding to the rows and sample corresponding to the columns therefore the simulated datasets are generated accordingly (**Table 1**). A random noise has been added for randomness of the data. We generated two types of datasets: i) Type A (small sample dataset): number of gene G=1000 and number of small, n=12 (n_1_=n_2_=6) and ii) Type B (large sample dataset): number of gene G=1000 and number of sample, n=60 (n_1_=n_2_=30). We considered 20 (2%) genes out of 1000 genes as differentially expressed genes (DEG) and they are separated into up-regulated (*P_1_*=10), and down-regulated (*P_2_*=10). Rest of the genes (980) was considered as equally expressed genes (EEG). The parameter values of *μ* and *σ^2^* was set to 0 and 0.1, respectively. The above set-up was used to generate 100 datasets from both types. The training and test datasets were constructed in a way such that the number of chosen random samples from normal and cancer groups were same. Our proposed algorithm “**IPLS-LDA**” was compared with the most popular classifiers such as SVM, LDA, naïve bayes, KNN, RF and PLS-LDA. The semantic flowchart of the proposed procedure has been presented in Fig.1.

**Figure 1.**
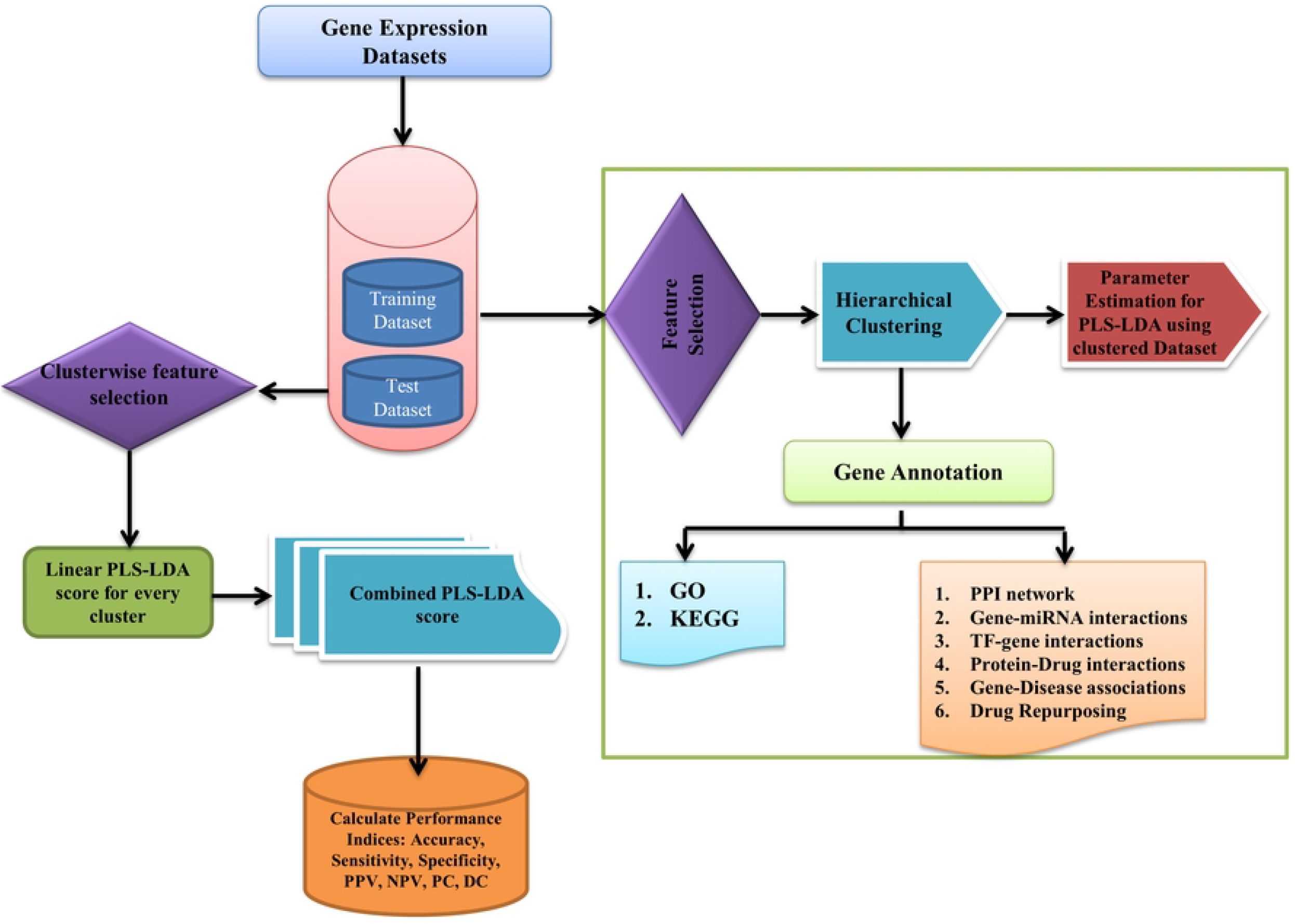
Semantic workflow of the proposed procedure.

**Table 1.**
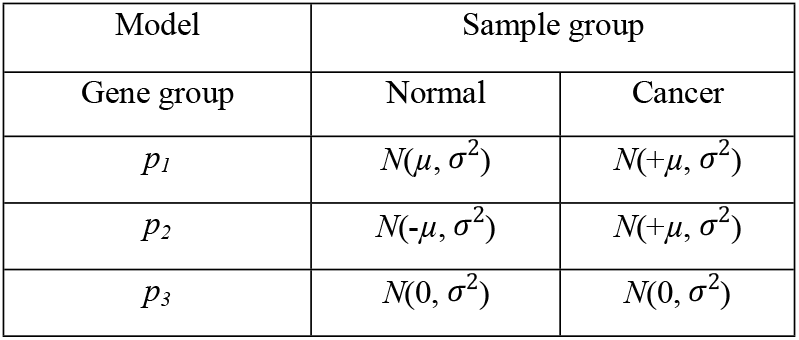
Simulated data generating model.

| Model | Sample group |  |
| --- | --- | --- |
| Gene group | Normal | Cancer |
| $p_1$ | $N(\mu, \sigma^2)$ | $N(+\mu, \sigma^2)$ |
| $p_2$ | $N(-\mu, \sigma^2)$ | $N(+\mu, \sigma^2)$ |
| $p_3$ | $N(0, \sigma^2)$ | $N(0, \sigma^2)$ |

To measure the classification performance of these seven segments, we calculated the nine different performance measures such as accuracy, upper and lower count, sensitivity, specificity, positive predictive value (PPV) and negative predictive value (NPV), PC and DC based on 20 DE genes for 100 datasets and the average values of these performance measures were recorded. These average values for both small and large sample cases were shown in Table 2 and Table 3, respectively. For IPLS-LDA we used hierarchical clustering to determine the number of gene clusters in GED. Fig. 2 (a) showed the clear picture of clusters using a dendrogram that divided the GED into two gene groups. To make it more clear we also applied average silhouette width to determine the number of cluster. This method also suggested that the optimal number of gene cluster was 2 (see Fig.2 (b)). It is noticed from Table 2 that, the IPLS-LDA algorithm outperformed the SVM, LDA, KNN, Naïve Bayes, RF and PLS-LDA classifiers for small sample case. For example Table 2 produces, the average accuracies of SVM, LDA, KNN, Naïve Bayes, RF, PLS-LDA and IPLS-LDA classifiers are 0.961, 0.938, 0.943, 0.960, 0.898, 0.964, and 0.970 respectively. Table 3 illustrates that, Naïve Bayes, PLS-LDA and IPLS-LDA produces almost similar performance over other four methods (for large sample case). For example, the average accuracies of SVM, LDA, KNN, Naïve Bayes, RF, PLS-LDA and IPLS-LDA classifiers are 0.975, 0.969, 0.970, 0.981, 0.931, 0.982 and 0.983 respectively. However, the IPLS-LDA produces slightly better performance than Naïve Bayes, and PLS-LDA.

**Figure 2.**
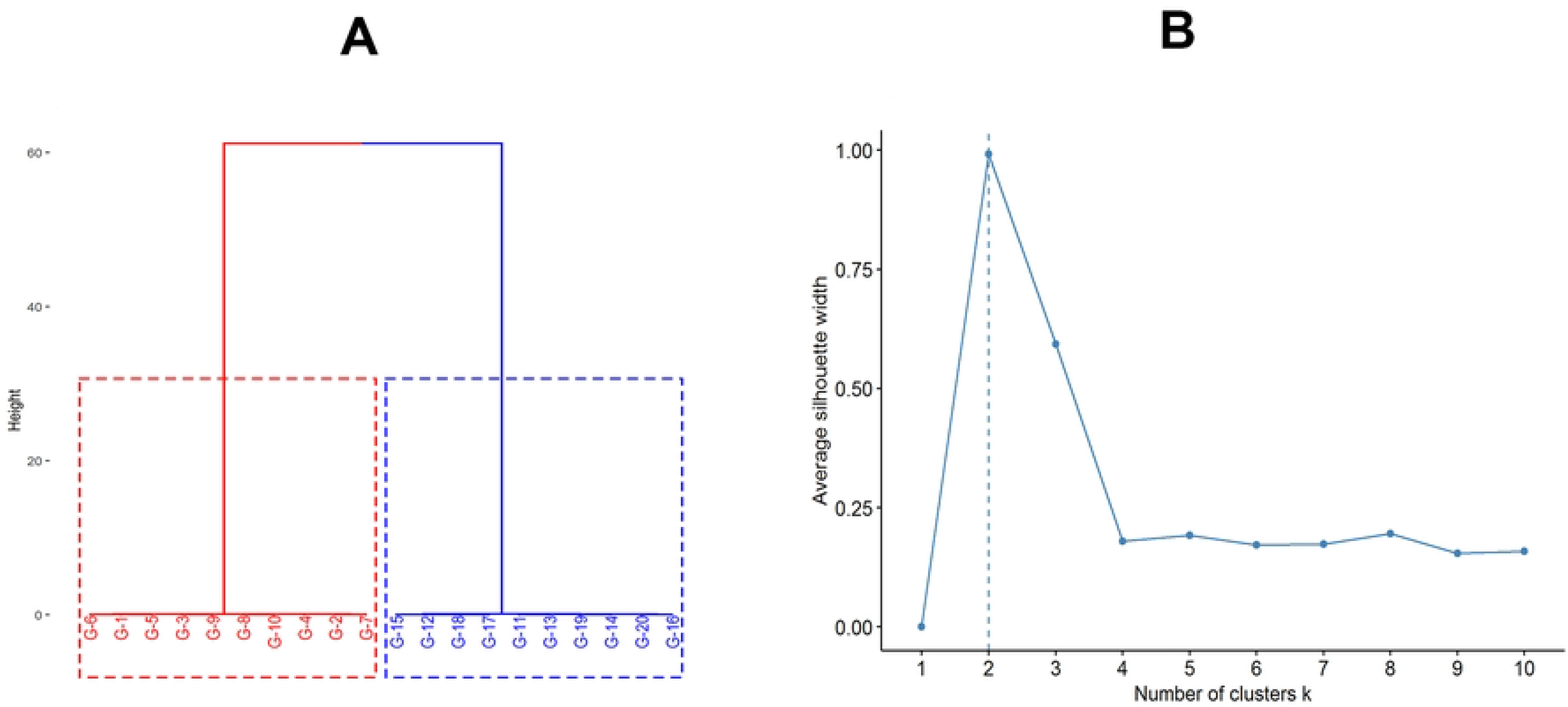
Optimal number of cluster determination. (A) cluster dendrogram, (B) plot of average silhouette width

The boxplot of test of accuracies for both small and large sample sizes were depicted in Fig. 3 (a) and Fig. 3 (b), respectively. Fig. S1 (a) and Fig. S1 (b) in the supplementary file, represent the ROC curve (FPR vs. TPR) for both the small sample and large sample size, respectively.

**Figure 3.**
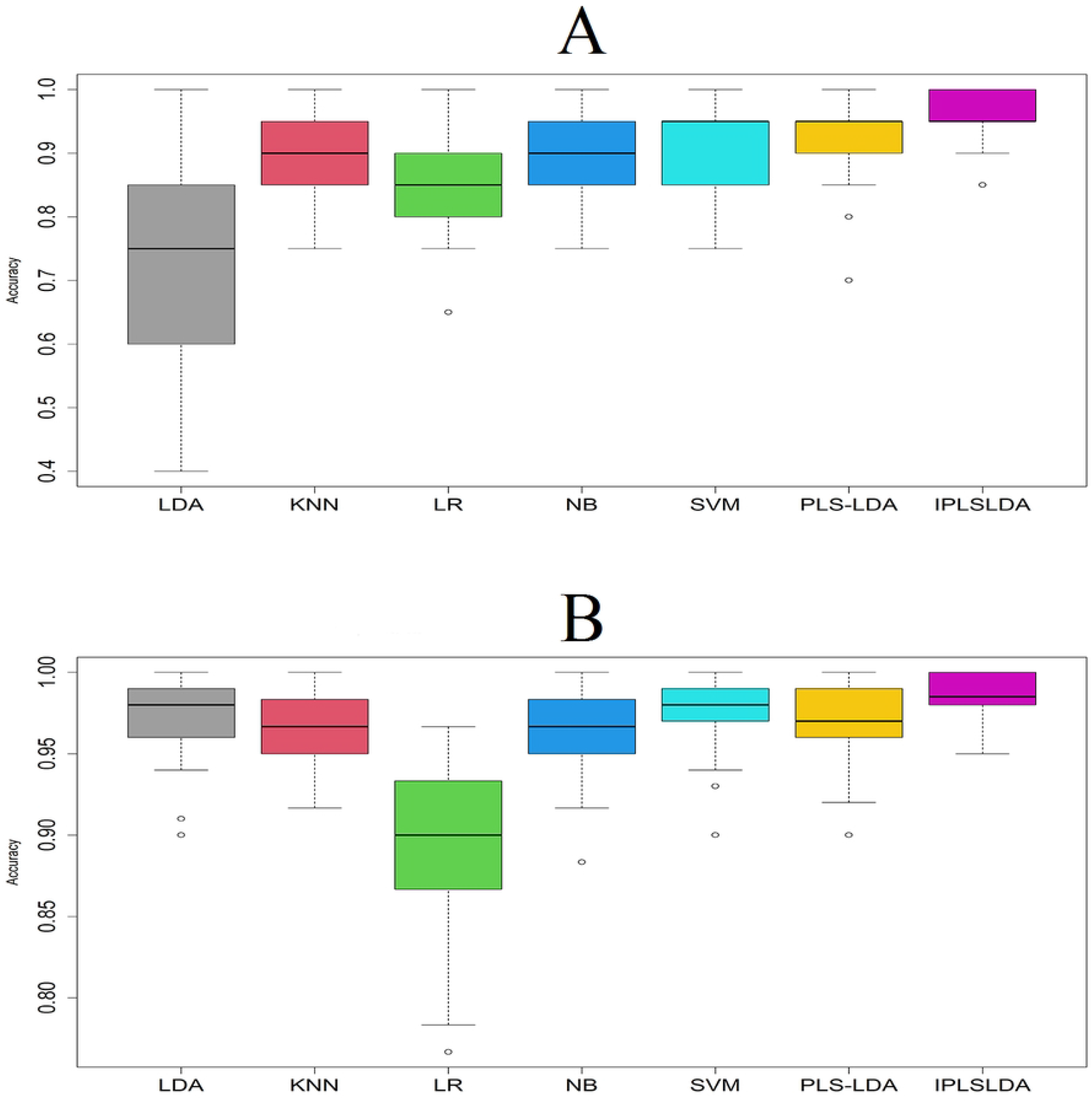
Performance evaluation of seven classifiers using boxplot of accuracies based on simulated datasets. (A) small-sample case, (B) large-sample case.

**Table 2.** Performance evaluation of the seven classifiers with small sample size (n_1_=n_2_=6) based on simulation data.

| Methods | SVM | LDA | KNN | NB | RF | PLS_LDA | IPLS-LDA |
| --- | --- | --- | --- | --- | --- | --- | --- |
| Accuracy | 0.961 | 0.938 | 0.943 | 0.960 | 0.898 | 0.964 | 0.970 |
| Lower Con.Int | 0.878 | 0.847 | 0.853 | 0.877 | 0.794 | 0.884 | 0.891 |
| Upper Con.Int | 0.991 | 0.980 | 0.984 | 0.991 | 0.959 | 0.992 | 0.994 |
| Sensitivity | 0.958 | 0.937 | 0.944 | 0.955 | 0.906 | 0.963 | 0.953 |
| Specificity | 0.963 | 0.938 | 0.942 | 0.964 | 0.890 | 0.966 | 0.986 |
| PPV | 0.964 | 0.94 | 0.944 | 0.965 | 0.896 | 0.967 | 0.986 |
| NPV | 0.960 | 0.939 | 0.946 | 0.957 | 0.906 | 0.964 | 0.956 |
| PC | 0.500 | 0.500 | 0.500 | 0.500 | 0.500 | 0.500 | 0.500 |
| DC | 0.479 | 0.469 | 0.472 | 0.478 | 0.453 | 0.963 | 0.953 |

**Table 3.** Performance evaluation of the seven classifiers with large sample size (n_1_=n_2_=30) based on simulation data.

| Methods | SVM | LDA | KNN | NB | RF | PLS_LDA | IPLS-LDA |
| --- | --- | --- | --- | --- | --- | --- | --- |
| Accuracy | 0.975 | 0.969 | 0.970 | 0.981 | 0.931 | 0.982 | 0.983 |
| Lower Con.Int | 0.925 | 0.916 | 0.916 | 0.932 | 0.864 | 0.934 | 0.936 |
| Upper Con.Int | 0.994 | 0.991 | 0.992 | 0.996 | 0.971 | 0.996 | 0.997 |
| Sensitivity | 0.974 | 0.970 | 0.976 | 0.978 | 0.935 | 0.980 | 0.975 |
| Specificity | 0.977 | 0.968 | 0.964 | 0.983 | 0.928 | 0.984 | 0.992 |
| PPV | 0.977 | 0.969 | 0.965 | 0.983 | 0.930 | 0.984 | 0.992 |
| NPV | 0.975 | 0.971 | 0.976 | 0.979 | 0.935 | 0.980 | 0.976 |
| PC | 0.500 | 0.500 | 0.500 | 0.500 | 0.500 | 0.500 | 0.500 |
| DC | 0.487 | 0.485 | 0.488 | 0.489 | 0.467 | 0.980 | 0.975 |

The results obtain from both boxplot and ROC curve also reflect the results of Table 2 and Table 3. Therefore, we may conclude that, IPLS-LDA produces better results than other six SML classification methods.

### 3.2. Real Data Analysis

#### 3.2.1. Colon Cancer data

The Colon cancer dataset was collected from a Colon cancer study [31] which consists of 6500 genes. 2000 of them were selected for the further analysis based on the minimal intensity of the samples. Therefore, the formation of Colon cancer dataset based on a data frame which corresponds to 2000 genes (rows) measured on 62 samples (columns). The first 40 columns from cancer samples and the rest of 22 columns are normal (healthy sample. This dataset can also be downloaded from plsgenomics [32] R package. At first we used a logarithmic transformation in the pre-processing step to reduce the effect of extreme values and make it normally distributed. To investigate the classification performance of the seven methods we constructed training and test datasets in a way such that 70% random samples belongs to training dataset and 30% random samples belongs to test dataset. To reconstruct a reduced training dataset, top 50 features were selected from training dataset using t-test by ranking the p value (<0.05). Now based on the reduced training dataset we evaluated the classification performance of seven methods and calculated the nine different performance indices such as accuracy, upper and lower count, sensitivity, specificity, positive predictive value (PPV) and negative predictive value (NPV), PC and DC. This process was repeated 100 times to record values of the various performance measures. The average values of these performance measurements showed in Table 4. For the IPLS-LDA, a hierarchical clustering method was used to determine the number of gene clusters in GED. Fig. 4 (a) showed the clear picture of clusters using a dendrogram that divided the GED into two gene groups. To make it more clear we also applied average silhouette width to determine the number of cluster. This method also suggested that the optimal number of gene cluster was 2 (see Fig.4 (b)). Table 4 describes that, SVM, KNN, PLS-LDA and IPLS-LDA produces better results than LDA, Naïve Bayes and RF. However, IPLS-LDA produces much better results than SVM, KNN and PLS-LDA. For example, the average accuracies of SVM, LDA, KNN, Naïve Bayes, RF, PLS-LDA and IPLS-LDA classifiers are 0.812, 0.773, 0.820, 0.772, 0.777, 0.831 and 0.841, respectively. The boxplot of test of accuracies for colon cancer dataset was represented in Fig. 4 (c) and the results obtain from boxplot also reflect the results of Table 4.

**Figure 4.**
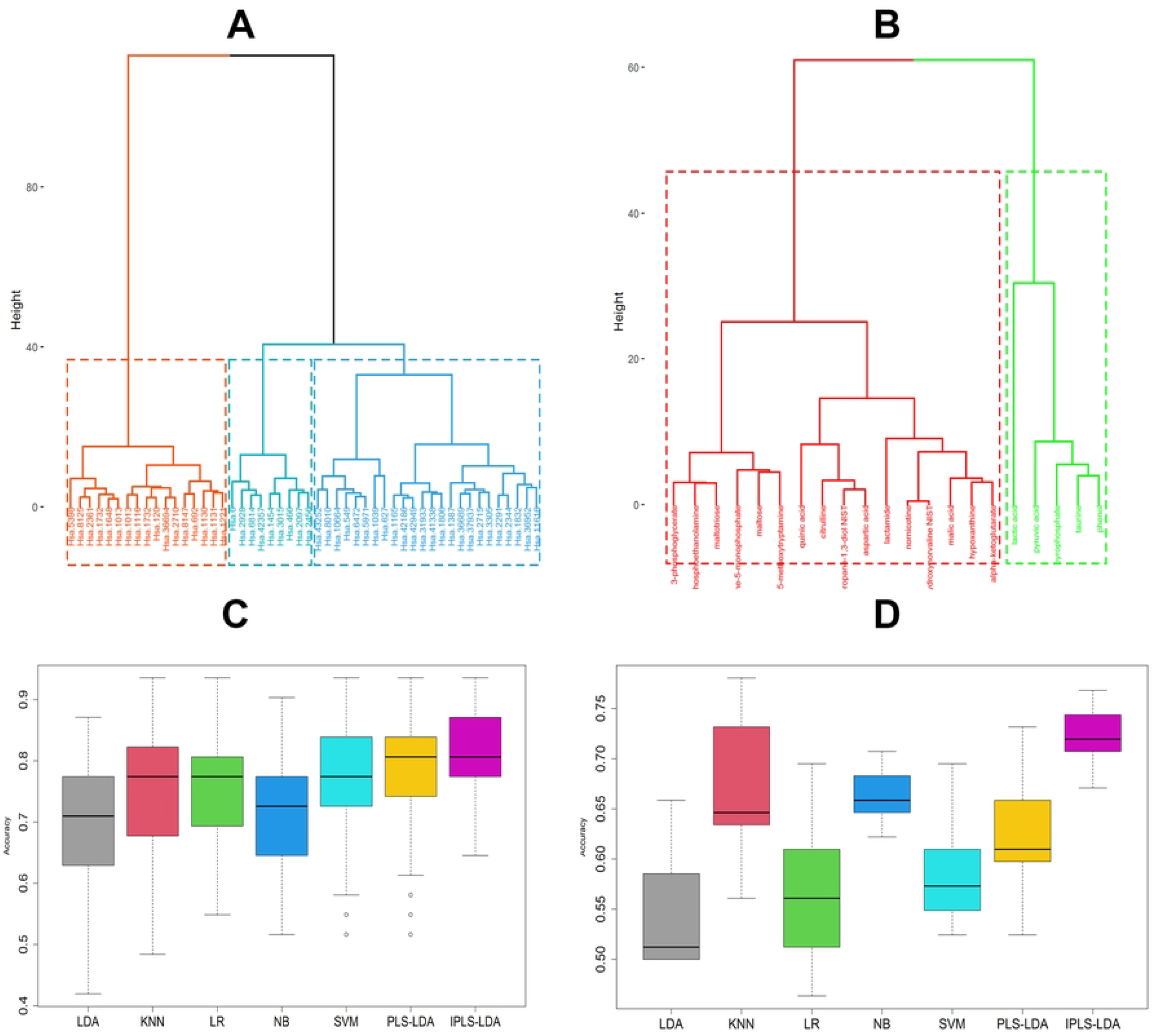
Performance evaluation of seven classifiers based on real datasets. (A) cluster dendrogram for colon cancer data, (B) cluster dendrogram for lung cancer metabolomics data,(C) boxplot of test accuracy for colon cancer data, (D) boxplot of test accuracy for lung cancer metabolomics data

**Table 4.**
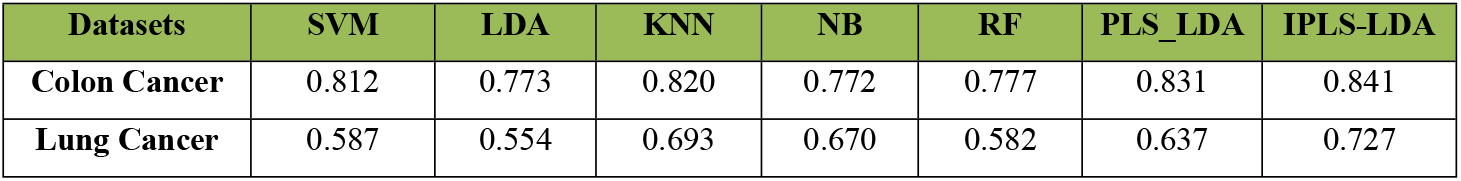
Performance comparison of the seven classifiers using Accuracy based on real datasets.

| Datasets | SVM | LDA | KNN | NB | RF | PLS_LDA | IPLS-LDA |
| --- | --- | --- | --- | --- | --- | --- | --- |
| Colon Cancer | 0.812 | 0.773 | 0.820 | 0.772 | 0.777 | 0.831 | 0.841 |
| Lung Cancer | 0.587 | 0.554 | 0.693 | 0.670 | 0.582 | 0.637 | 0.727 |

Genes a together with an interactive a network. To decode the biological mechanisms of top 50 DEGs (28 from cluster 1 and 22 from cluster 2) identified by the proposed procedure we revealed different gene ontology (GO) categories such as biological process (BP), cellular component (CC), and molecular functions (MF). The GO results were summarized in Supplementary Table. A1. We extracted the Kyoto Encyclopedia of Genes and Genomes (KEGG) pathway annotations of two cluster DEGs using Database for Annotation, Visualization, and Integrated Discovery (DAVID) [33]. The KEGG pathway enrichment analysis results have been summarized in Table.5. 28 genes of cluster 1 are enriched in different pathways (Ribosome, Hepatitis B, Spliceosome, IL-17 signaling pathway HTLV-I infection etc.) and 22 genes of cluster 2 are enriched in pathways (Thyroid hormone signaling pathway, Transcriptional misregulation in cancer, Pathways in cancer, Colorectal cancer etc.)

**Table 5.** Top 10 significantly enriched KEGG pathways for 50 DEGs identified by the proposed method in each clusters.

| Pathway | Total | Expected | Hits | P.Value |
| --- | --- | --- | --- | --- |
| Ribosome | 153 | 11.2 | 71 | 8.03E-41 |
| Hepatitis B | 163 | 11.9 | 43 | 4.21E-14 |
| Spliceosome | 134 | 9.82 | 38 | 1.11E-13 |
| Epstein-Barr virus infection | 201 | 14.7 | 47 | 3.53E-13 |
| Viral carcinogenesis | 201 | 14.7 | 47 | 3.53E-13 |
| Measles | 138 | 10.1 | 36 | 7.63E-12 |
| Basal transcription factors | 45 | 3.3 | 20 | 8.06E-12 |
| IL-17 signaling pathway | 93 | 6.82 | 27 | 2.44E-10 |
| Hepatitis C | 155 | 11.4 | 36 | 2.78E-10 |
| HTLV-I infection | 219 | 16 | 44 | 4.40E-10 |
| Pathway | Total | Expected | Hits | P.Value |
| Spliceosome | 134 | 5.92 | 33 | 1.88E-16 |
| Proteasome | 45 | 1.99 | 20 | 5.42E-16 |
| Thyroid hormone signaling pathway | 116 | 5.13 | 30 | 1.11E-15 |
| RNA transport | 165 | 7.29 | 33 | 1.24E-13 |
| Transcriptional misregulation in cancer | 186 | 8.22 | 32 | 2.13E-11 |
| Epstein-Barr virus infection | 201 | 8.88 | 33 | 3.78E-11 |
| Viral carcinogenesis | 201 | 8.88 | 33 | 3.78E-11 |
| HTLV-I infection | 219 | 9.68 | 34 | 9.16E-11 |
| Pathways in cancer | 530 | 23.4 | 56 | 4.71E-10 |
| Colorectal cancer | 86 | 3.8 | 20 | 5.93E-10 |

In addition, a protein-protein interaction (PPI), DEGs-miRNA, TFs-DEGs networks were also constructed using IMEx Interactome, miRTarBase and JASPER databases [34–36] via online tool NetworkAnalyst [37]. These networks were visualized in Fig.5 and Fig.S2, respectively. Therefore in colon cancer study, the IPLS-LDA produces better results than other methods.

**Figure 5.**
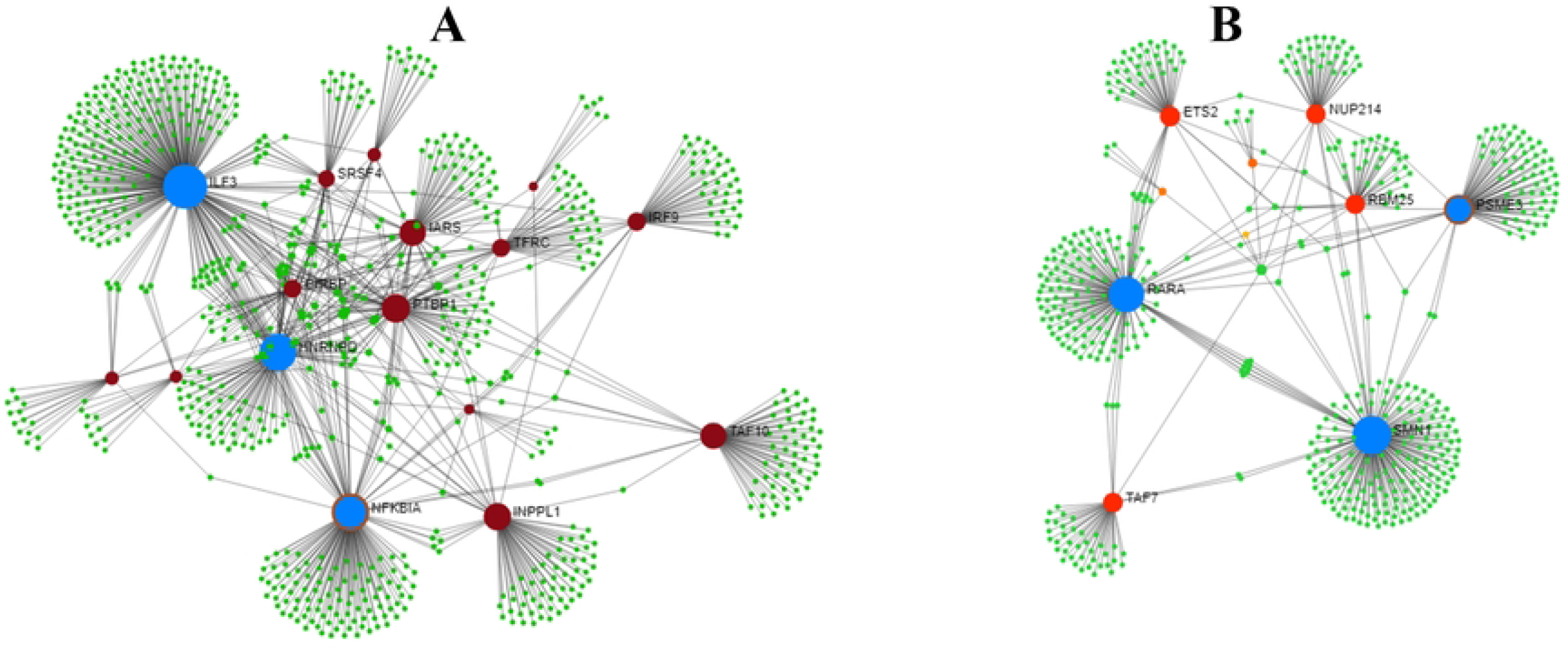
Protein-protein interaction (PPI) network analysis of 50 DEGs identified by proposed method using colon cancer data. (A) PPI network for cluster 1 (28 DEGs), (B) PPI network for cluster 2 (22 DEGs),

#### 3.2.2. Lung Cancer Metabolomics Data

The lung cancer metabolic dataset contain 158 metabolites. The data consists of total 82 blood samples (41 control and 41 cancer) produced by Oliver Fiehn [38] and these blood samples were collected by EDTA tubes (stored at −80 °C) and approved protocols were used to make over the samples (serum and plasma). Both the training (serum) and test (plasma) datasets consist of 158 metabolites with 82 samples. Then a pre-processing step was conducted by using a logarithmic transformation to reduce the effect of extreme values. For further making a reduced training datasets, top 50 feature was selected using t-test by ordering p-value (<0.05). To measure the classification performance of these seven methods, we calculated accuracy based on top 50 feature and the average values of these performance measures were recorded which shown in Table 5. For the IPLS-LDA, we used hierarchical clustering to determine the number of gene clusters in metabolic datasets. Fig. S3 (a) showed the clear picture of clusters using a dendrogram that divided the metabolic datasets into two gene groups. To make it more clear we also applied average silhouette width to determine the number of cluster. This method also suggested that the optimal number of gene cluster was 2 (see Fig.S3 (b)). It is noticed from Table 5 that, the IPLS-LDA algorithm outperformed the SVM, LDA, KNN, Naïve Bayes, RF and PLS-LDA classifiers for metabolic datasets. For example Table 5 produces, the average accuracies of SVM, LDA, KNN, Naïve Bayes, RF, PLS-LDA and IPLS-LDA classifiers are 0.587, 0.554, 0.693, 0.670, 0.582, 0.637 and 0.727 respectively. The boxplot of test of accuracies was depicted in Fig. 4 (d). The results obtain from boxplot also reflected in the results of Table 5. Therefore in metabolic data analysis, the IPLS-LDA produces better results than other methods.

## 4. Conclusions

In Bioinformatics, class prediction is one of the significant issues especially for microarray gene expression and metabolic datasets. Large number of data contained curse of dimensionality, non-linearity with heterogeneity that makes the analysis way difficult. Partial Least Square Linear Discriminant Analysis (PLS-LDA) is extensively used in gene expression (GE) and metabolomics datasets for predicting unclassified test data. However, this method also suffers from the problems discussed above. Hence, in this study, an improved PLS-LDA (IPLS-LDA) was developed by capturing the heterogeneity of features through an unsupervised hierarchical clustering approach. We investigated the performance of IPLS-LDA in a comparison of six popular classifiers though feature selection based on simulated, one gene expression and one metabolomics data analysis. Both simulated and real data analysis confirmed the superiority of the proposed algorithm than the other SML approaches in terms of various performance indices.

## Data availability

The proposed algorithm has been implemented in an R package, Uplsda was given in the https://github.com/snotjanu/UplsLda

---

**Figure S1. Performance evaluation of seven classifiers using ROC curve based on simulated datasets.** (A) small-sample case, (B) large-sample case.

**Figure S2. DEGs-miRNA and TFs-DEGs interaction network for colon cancer dataset.** (A) DEGs-miRNA for cluster 1, (B) DEGs-miRNA for cluster 2, (C) TFs-DEGs for cluster 1, (D) TFs-DEGs for cluster 2.

